# Predicting the Genomic Resolution of Bulk Segregant Analysis

**DOI:** 10.1101/2021.04.16.440220

**Authors:** Runxi Shen, Philipp W. Messer

## Abstract

Bulk segregant analysis (BSA) is a technique for identifying the genetic loci that underlie phenotypic trait differences. The basic approach of this method is to compare two pools of individuals from the opposing tails of the phenotypic distribution, sampled from an interbred population. Each pool is sequenced and scanned for alleles that show divergent frequencies between the pools, indicating potential association with the observed trait differences. BSA has already been successfully applied to the mapping of various quantitative trait loci in organisms ranging from yeast to maize. However, these studies have typically suffered from rather low mapping resolution, and we still lack a detailed understanding of how this resolution is affected by experimental parameters. Here, we use coalescence theory to calculate the expected genomic resolution of BSA. We first show that in an idealized interbreeding population of infinite size, the expected length of the mapped region is inversely proportional to the recombination rate, the number of generations of interbreeding, and the number of genomes sampled, as intuitively expected. In a finite population, coalescence events in the genealogy of the sample reduce the number of potentially informative recombination events during interbreeding, thereby increasing the length of the mapped region. This is incorporated into our theory by an effective population size parameter that specifies the pairwise coalescence rate of the interbreeding population. The mapping resolution predicted by our theory closely matches numerical simulations. Furthermore, we show that the approach can easily be extended to modifications of the crossing scheme. Our framework enables researchers to predict the expected power of their mapping experiments, and to evaluate how their experimental design could be tuned to optimize mapping resolution.

## Introduction

The advent of easy and affordable genome sequencing has enabled powerful genetic mapping approaches. In addition to improving our understanding of the molecular basis of phenotypic traits, such approaches can have important practical applications. For example, genetic mapping can help us identify variants that underlie human diseases [1], localize genes associated with favorable traits in plant or animal breeding [10, 37], and detect the loci responsible for drug or pesticide resistance [3, 28, 29].

Various techniques have been developed for this purpose, ranging from classical linkage mapping to genome-wide association studies (GWAS), with numerous extensions or combinations of these approaches that are often tailored towards specific applications. Which particular technique is best suited for a given problem can depend on a variety of factors, such as the genetic architecture of the trait, the specific biology of the study system, the resources available for experiments and sequencing, and the mapping resolution desired.

In species that can be experimentally crossed, classical linkage mapping has proven a powerful technique for detecting quantitative trait loci (QTL) [20, 22, 25, 39]. This method involves the generation of an F_1_ cross from two parental homozygous strains of contrasting phenotypes. The F_1_ offspring are then backcrossed to the parental strains, and the resulting progeny are phenotyped for the trait of interest and genotyped at a set of marker loci distributed across the genome. By scanning for markers with an inheritance pattern that correlates with the trait, one can localize the segments of the genome on which causal variants could reside. This method has long been the primary genetic mapping technique. However, it tends to attain rather low genomic resolution (i.e., the length of the identified genomic region in which the causal locus must be contained but cannot be more precisely pinpointed). This is because the segments linked to the parental strains are typically quite long due to the limited number of recombination events in a single cross.

GWAS is an alternative approach for QTL mapping that does not require experimental crosses, but instead exploits historical recombination events in a genetically diverse population. In this approach, a large number of individuals are genotyped at a dense set of SNP markers, or by whole genome sequencing, and phenotyped for the trait of interest [34]. The QTL responsible for trait variation can then be identified by regressing SNP genotypes against phenotype. The genomic resolution of this approach is limited in principle only by the density of SNP markers and the genomic distance over which linkage disequilibrium decays in the mapped population. As a result, GWAS can sometimes detect even individual causal SNPs and tends to work well also for highly polygenic traits, whereas classical linkage mapping is more suited for traits with a few major QTL. However, the trait of interest needs to exhibit sufficiently high levels of additive genetic variation in the population for GWAS to work. Detection power also tends to be limited for causal variants that segregate at low population frequency. In addition, due to the large number of SNPs tested, the thresholds for calling statistical significance can be very high.

Bulk segregant analysis (BSA) is a mapping approach that combines ideas from linkage mapping and GWAS [24]. Like classical linkage mapping, BSA starts from two homozygous parental strains of contrasting phenotypes. These strains are then crossed to generate an F_1_ population that is further interbred for several generations while maintaining a sufficiently large population size to allow recombination to break up linkage from the two parental strains. In the final generation, two pools of individuals are selected from the tails of the phenotypic distribution, and each of these pools is sequenced. The alleles responsible for trait differences (as well as any alleles linked to them) should then exhibit significant frequency differences between the two pools, while alleles at other loci should be present in both pools at similar frequencies.

In contrast to both GWAS and classical linkage mapping, BSA does not require the sequencing of individuals, since only the overall allele frequencies in the two pools are relevant. This allows the use of more economic sequencing approaches such as Pool-seq [30]. The resolution of BSA is still expected to be considerably lower than GWAS because the number of generations over which the population is interbred will be limited. For longer experiments, the effects of drift could also become problematic [26]. However, in contrast to GWAS, BSA can still be used for detecting QTL where causal alleles are segregating at low frequency in the population, as long as they are present in one of the parental strains. This could be a key factor for applications such as the mapping of drug or pesticide resistance mutations.

Conceptually, BSA is similar to “introgression mapping” [7, 31], where the population is repeatedly selected for the phenotype of the first parental strain in every even generation of the experiment. The surviving individuals are then back-crossed to the second parental strain and the resulting offspring are interbred without selection in every odd generation. Under this approach, the population at the end of the experiment should be genetically similar to the first strain in the genomic regions that surround causal QTL, while it should be similar to the second parental strain for the rest of the genome. Note, however, that this approach can require a considerably higher experimental effort than BSA.

BSA has already been successfully applied in various contexts. For example, implementations of this approach have been used to identify DNA markers linked to disease-resistance genes in lettuce [24] and pest-resistance genes in crops [32], to study horizontal gene transfer in *Tetraychus urticea* [4], to locate QTLs associated with drought resistance in maize [27], and to map the genetic basis of various complex traits in yeast and *Drosophila* [8, 18, 21].

Despite these successful applications, one practical shortcoming of BSA is its tendency to produce very wide peaks of significance, which in previous studies have often extended over hundreds of kilobases [38] or even several megabases [33]. This is particularly problematic because we do not currently have a good understanding of how the expected mapping resolution is determined by biological and experimental parameters. Simulation studies have shed some light on this issue and demonstrated that more generations of interbreeding, a larger population size during interbreeding, and deeper sequencing can all improve mapping resolution, while the size of the selected pools apparently has less of an impact [26]. Nevertheless, it would still be desirable to have an analytical understanding of exactly how all of these factors influence mapping resolution; this would allow us to predict the expected resolution for a given experiment, and to assess which factors one should tune to optimize the mapping resolution most economically.

In this study, we employ coalescence theory to develop an analytical framework for calculating the expected mapping resolution of a BSA experiment. Our theory reveals how the recombination rate of the study organism, the effective population size during interbreeding, the overall length of the experiment, and the number of genotyped individuals combine to determine the maximally achievable mapping resolution for a trait with a simple genetic architecture.

## Results

Consider a phenotypic trait determined by a single QTL with two segregating alleles: *A* and *a*. We assume that recombination occurs at a uniform rate *r* per bp along the chromosome. For simplicity, we neglect gene conversion and assume that recombination events always result in crossover. Starting from the two founding strains (“blue” and “red”) which we assume have genotypes *AA* and *aa*, respectively, a BSA experiment is performed for *t* generations of interbreeding, as outlined in Figure 1A. At the end of the experiment, we select two samples from the interbred population, such that the first sample contains *s* chromosomes of genotype *A*, while the second contains *s* chromosomes of genotype *a*. In practice, this could be achieved be selecting *s*/2 individuals that are homozygous for *A* as the first sample, and *s*/2 individuals that are homozygous for *a* as the second sample (our approach thus relies on the ability to accurately identify such individuals based on their phenotype). Each of the two samples is then pooled and sequenced.

**Figure 1:**
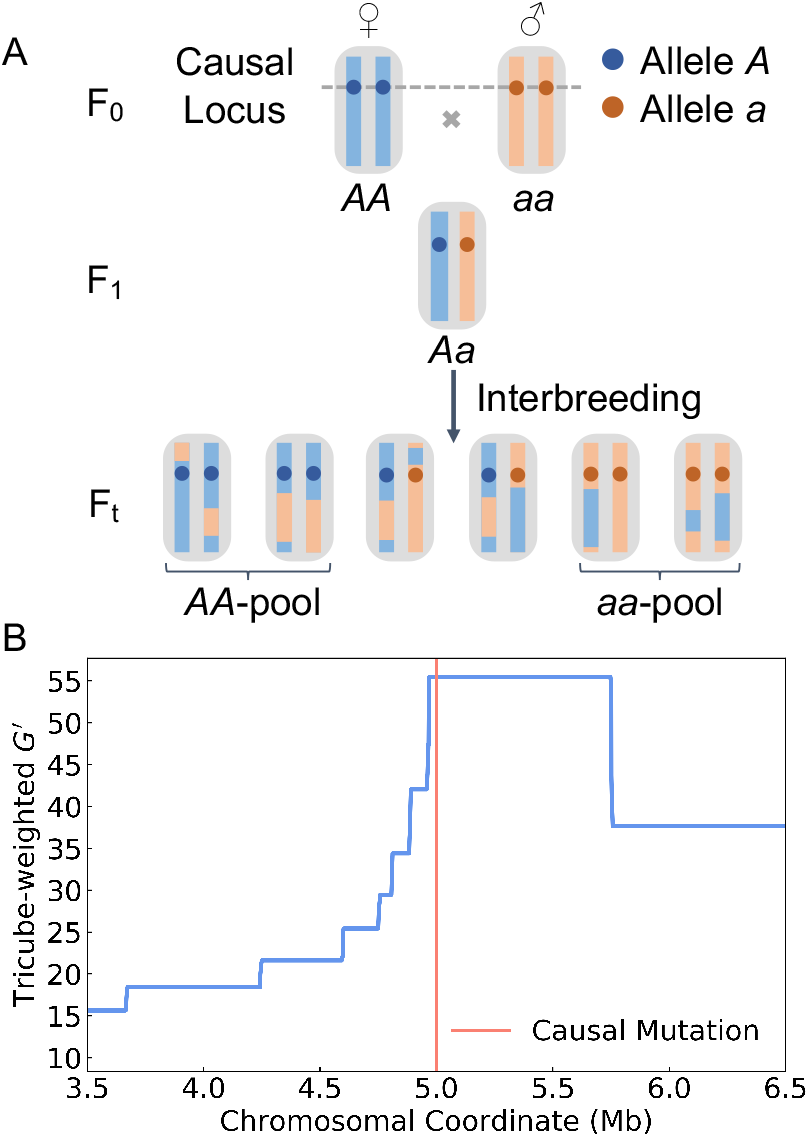
Illustration of BSA. (A) Our model assumes a trait determined by a single QTL with two different alleles (*A, a*). The starting point of the experiment are two homozygous parental strains, represented by red and blue chromosomes. The blue strain carries the *A* allele and the red carries the *a* allele. An F_1_ population is created and interbred for *t* generations. At the end of the experiment, two pools of individuals are selected such that the first comprises only *AA* individuals and the second only *aa* individuals. The mapping resolution is determined by the length of the region surrounding the QTL for which all chromosomes in the *AA*-pool still have blue ancestry, while all in the *aa*-pool still have red ancestry. (B) Mapping resolution in a simulated BSA experiment for a QTL located at the center (red line) of a 10 Mb-long chromosome. Interbreeding was modeled for 10 generations in a population of 100 individuals with a uniform recombination rate of 1.0 cM/Mb (see Methods). Two pools of 10 *AA* and 10 *aa* individuals were selected at the end of the experiment. The blue curve shows the *G*′ statistic estimated from marker SNPs placed at equidistant intervals of 1 kb along the chromosome to differentiate ancestry from the two parental strains. The peak in *G*′ around the QTL indicates the region where all chromosomes in the *AA*/*aa* pools still have blue/red ancestry, which extends for ∼ 0.5 Mb.

As a consequence of recombination during interbreeding, each of the chromosomes sampled at the end of the experiment should be a mosaic of red and blue ancestry segments. However, there should be a region surrounding the QTL where all chromosomes in the *AA*-pool still have blue ancestry, while all in the *aa*-pool still have red ancestry. The maximally achievable mapping resolution is determined by the size of this region (assuming that there is only one such region in the sample). Several summary statistics have been developed to identify such regions, which typically rely on the detection of differences in allele frequencies at marker SNPs between the two pools. Examples for such statistics include ancestry difference (*A*_*d*_) [26], Δ(SNP −index) [9], and a modified G-statistic (*G*′) [21]. An illustration of this mapping problem is provided in Figure 1B, where we show *G*′ estimated along a chromosome in a simulated BSA experiment.

The goal of our theoretical analysis will be to calculate the expected length of the region where all chromosomes in the *AA*-pool still have blue ancestry, while all in the *aa*-pool still have red ancestry. For this purpose, let us define *D* as the distance to the closest “ancestry breakpoint” (defining a point where ancestry changes between blue and red in a chromosome) located downstream of the QTL among all chromosomes in the samples (Figure 2A). Due to symmetry, the expected length of the mapped region will then be simply 2*E*[*D*], where *E*[*D*] denotes the expectation value of *D* (we will neglect edge effects when a QTL is located close to the start/end of the chromosome). This length determines the expected mapping resolution of the BSA experiment (with “shorter” expected mapping tract lengths corresponding to “higher” resolution). Note that the actually achievable resolution will likely be lower in practice than predicted by our theory due to the need to rely on marker SNPs as proxies for ancestry, as well as other experimental factors such as sequencing errors. Our general approach for the calculation of *E*[*D*] is to trace the lineages of all sampled chromosomes back to the two parental strains, and then study how ancestry breakpoints have been generated along this genealogy (Figure 2B). Note that due to recombination events, local genealogies will vary as one moves along the chromosome of the samples, constituting the so-called “tree sequence” [17]. However, at any given position, there will be exactly one genealogy. Thus, the lineage of any given sampled chromosome at that position can be traced back all the way to a single chromosome in the F_0_. If this happens to be a red chromosome, the sampled chromosome will be assigned red ancestry at this position, otherwise it will be assigned blue ancestry.

**Figure 2:**
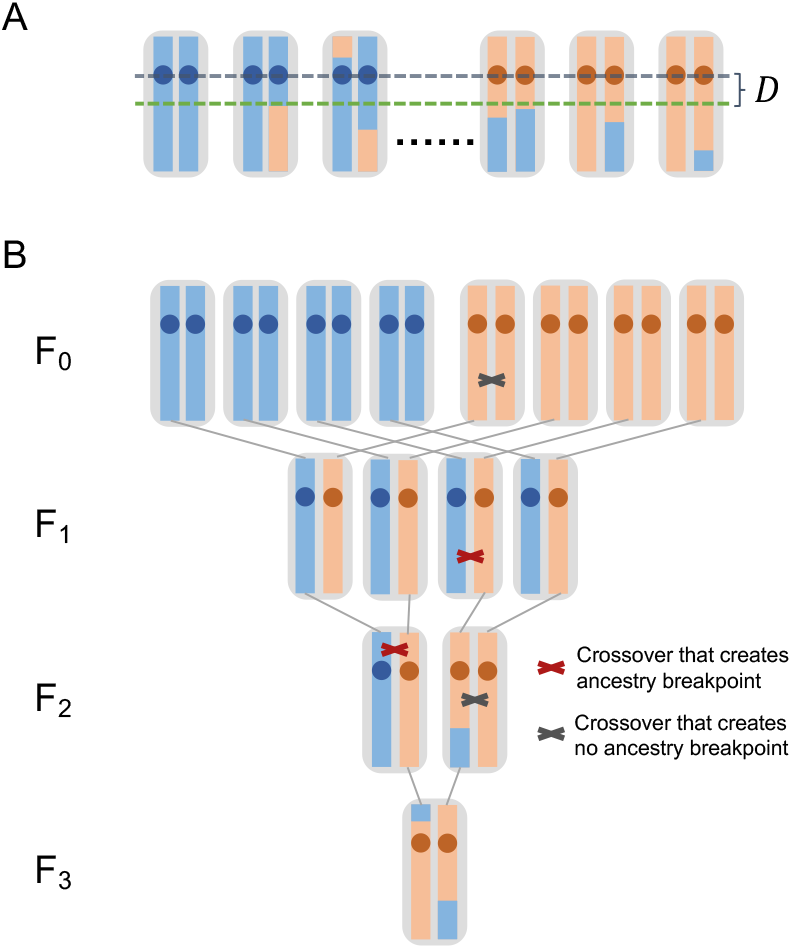
Resolution of a BSA experiment. (A) We define *D* as the distance between the QTL and the first ancestry breakpoint downstream of the QTL in all samples. (B) Example of a full pedigree of an individual from the F_3_. All crossover events that have occurred along its pedigree are also shown. Only those crossover events occurring in individuals that carried red and blue ancestry at the location of the crossover actually generated new ancestry breakpoints, and every breakpoint observed in the sampled chromosome can be traced back to such a specific crossover event in the pedigree.

Each ancestry breakpoint in a sampled chromosome stems from a crossover event in one of its ancestors. Importantly, this must have been an ancestor that carried a blue ancestry segment around the crossover location in one of its chromosomes, and a red one in the other (Figure 2B). By contrast, crossover events at positions where an individual carries either two blue or two red ancestry segments around the crossover location will never create new ancestry breakpoints.

### Infinite Population Model

We initially want to assume an idealized model of an interbreeding population of infinite size. This is for two reasons: first, we want to be able to neglect coalescence events when tracing back the lineages from the chromosomes in our sample to the chromosomes in parental strains. Second, we want to be able to neglect any changes in allele frequencies over the course of the experiment due to random genetic drift.

Let us first consider a short BSA experiment where the population is already sampled in the F_2_. Since all individuals in the F_1_ carry one red and one blue chromosome, all recombination events in this generation should create new ancestry breakpoints. We model a uniform recombination rate (*r*) per base pair. In 2*s* sampled chromosomes from the F_2_ (representing the combined two pools), the overall rate (*R*) at which new ancestry breakpoints have been created per bp in this generation is therefore simply *R* = *r* × 2*s*.

Assuming *R* ≪ 1, we can model these events by a Poisson process along the chromosome. The distance *D* to the closest ancestry breakpoint downstream of the QTL in all sampled chromosomes should then be an exponential random variable with cumulative density function *P*(*D* ≤ *d*) = 1 − *e*^−*Rd*^ and expectation value *E*[*D*] = 1/*R* = 1/(2*rs*).

We can directly extend this process to chromosomes sampled from the F_3_, but here things become a bit more complicated. This is because the parents of the sampled individuals are no longer guaranteed to carry one red and one blue chromosome. Instead, according to Hardy-Weinberg equilibrium, the probability that a randomly picked individual from the F_2_ at any given genomic position will carry chromosomes with different ancestry is only 1*/*2. Thus, only half of the crossover events during meiosis are actually expected to create new ancestry breakpoints in this generation, and the overall rate at which new ancestry breakpoints are created per bp is therefore *R* = *rs*. Since we neglect drift in the infinite population model, this should be the same fraction for all future generations.

The infinite population model also ensures that no two sampled chromosomes will ever share a parent or grandparent with each other. Consequently, we can model individual ancestral lineages completely independently of each other. In 2*s* chromosomes sampled from the F_3_, the overall rate (*R*) at which new ancestry breakpoints have been generated per bp is therefore simply the sum of the individual rates over the 2*s* lineages and the two parental generations: *R* = 2*rs* + *rs* = 3*rs*. Every additional generation of crossing will further increment this rate by *rs*. Thus, after *t* generations of interbreeding, the overall rate will be *R* = *rst*. Assuming *R* ≪ 1, we can again model these events by a Poisson process along the chromosome, yielding:

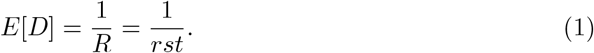

Because the situation upstream and downstream of the QTL is symmetric, the expected resolution of the BSA experiment in this infinite population mode is then simply 2*E*[*D*] = 2*/*(*rst*) bp. Thus, it is inversely proportional to the product of the recombination rate, sample size, and length of the experiment. This result is very intuitive; all that matters is the overall rate at which new ancestry breakpoints are generated along the lineages of the sample. Note that because we have modeled *D* as an exponential random variable, its variance will be given by 1*/*(*rst*)^2^ and the full cumulative distribution function will be *P* (*D* ≤ *d*) = 1 − *e*^*−rstd*^.

### Finite Population Model

In the infinite population model, every ancestry breakpoint present in the sampled chromosomes traces back to a unique crossover event along the genealogy of the sample. In a finite population, different chromosomes can share a breakpoint that traces back to the same crossover event in a common ancestor. In that case, we can no longer describe the genealogy of the samples by 2*s* distinct lineages through the interbreeding phase, since individual lineages could have merged (Figure 3).

**Figure 3:**
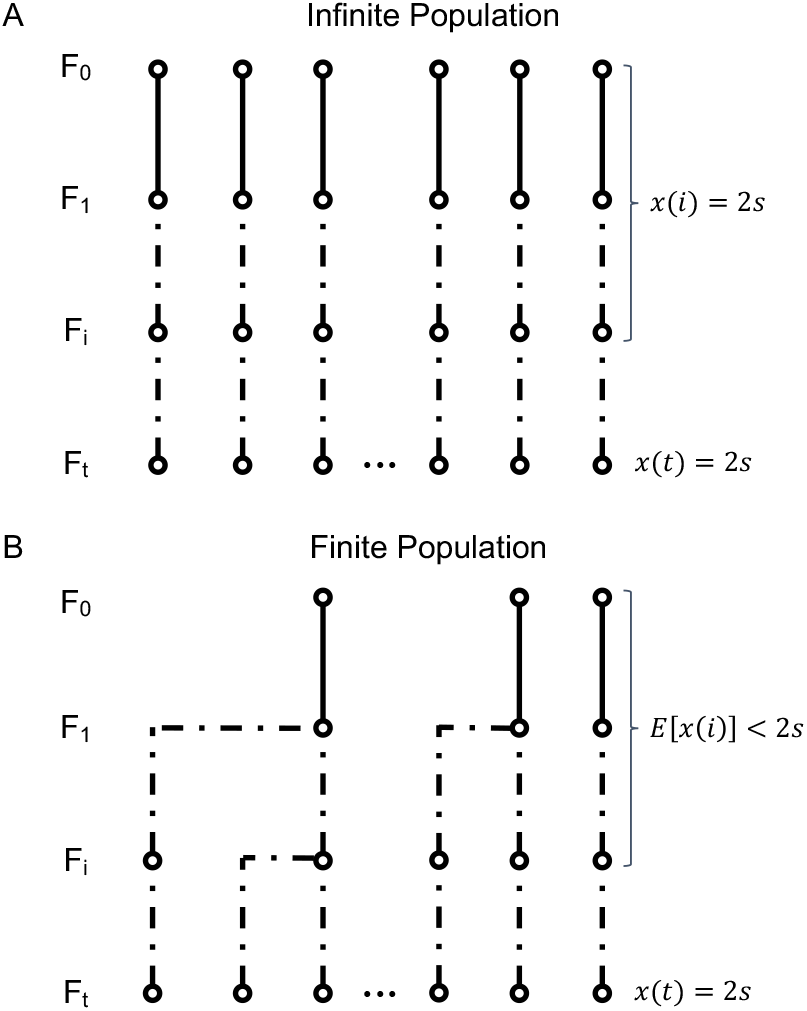
Infinite and finite population models. (A) In the infinite population model, all lineages descend independently and the overall length of the genealogy of 2*s* sampled genomes is simply 2*st*. (B) In the finite population model, by contrast, lineages can coalesce in ancestors of the sample, reducing the expected overall number of ancestors in previous generations and thereby the expected length of the genealogy.

An important consequence of this is that the expected “length” of the sample’s genealogy will be shorter in a finite population compared to our infinite population model, where it was simply 2*st*. In general, this should reduce the number of ancestry breakpoints captured in the sample, thereby increasing the length of the mapped region.

To derive an analytic expression for the mapping resolution in a finite population, let us assume that we can model it as a diploid Wright-Fisher population with coalescence effective population size *N*_*e*_. Let *x*(*i*) denote the number of ancestral lineages in the sample’s genealogy in generation *i* at a given genomic position (Figure 3). We can calculate how *x*(*i*) is expected to change between consecutive generations, applying a result from the theory of occupancy distributions [15]:

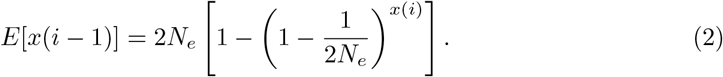

Evaluating this equation recursively, starting from *x*(*t*) = 2*s*, then allows us to calculate the expectation values of *x*(*i*) all the way back to *i* = 2.

As in the infinite population model, every crossover event in the F_1_ will create a new ancestry breakpoint, while this should be true for only half of such events in subsequent generations. Together with the above result for *x*(*i*), this allows us to calculate the overall rate (*R*) at which new ancestry breakpoints are generated per bp along the genealogy of all sampled chromosomes:

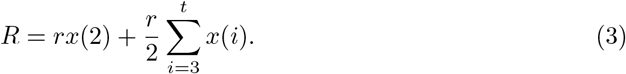

Since *E*[*x*(*i*)] < 2*s* for all *i* < *t*, this rate will be smaller than the corresponding rate *R* = *rst* of the infinite population model.

One important assumption underlying Eq. (3) is that the population frequencies of red and blue alleles still remain constant at 50% over the course of the experiment, so that from the F_2_ onward, the probability that a randomly chosen individual carries both a red and a blue ancestry segment at any given genomic position remains at 0.5. However, random genetic drift should lead to a decay of heterozygosity (*H*) over time according to *H* ∝ exp(−*t*/(4*N*_*e*_)], and the probability that an individual carries ancestry segments from both parental strains at a given genomic position is expected to decrease at a similar rate. Since we neglect this effect, Eq. (3) should still overestimate *R*, although much less so than in the infinite population model. This should primarily be a problem for very long BSA experiments with small *N*_*e*_ where *t* ≪ 4*N*_*e*_ does not hold.

As long as *R* ≪ 1, we can again model the creation of new ancestry breakpoints by a Poisson process along the chromosome. The distance *D* to the closest ancestry breakpoint downstream of the QTL captured in the sample will then be an exponential random variable with expectation value:

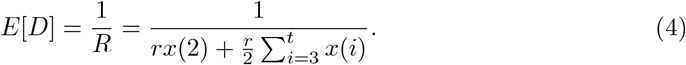

This result provides an analytic solution for the expected mapping resolution of a BSA experiment with an interbreeding population of effective size *N*_*e*_. However, its calculation requires iterative evaluation of Eq. (2), and we are not aware of any closed-form solution for this recursion. Even though all elements of *x*(*i*) can be easily calculated with the help of a computer, this may not be particularly helpful in allowing us to understand how individual parameters are expected to affect the mapping resolution. To address this issue, we will make use of a previously suggested deterministic approximation for *x*(*i*), which can be obtained by mapping the recursion to a differential equation [6, 11, 14, 23]:

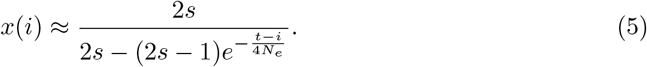

We will further replace the summation in Eq. (3) by an integral over the *t* generations of the experiment, yielding:

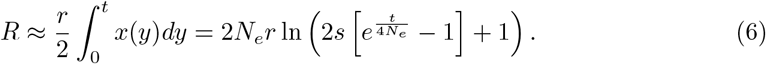

Note that this integration assumes that recombination events along the genealogy create ancestry breakpoints with a uniform probability of 1*/*2 in every generation (not just from the F_2_ onward). This assumption is obviously incorrect for individuals in the F_1_, where every recombination event will generate a new ancestry breakpoint. However, by extending our integration back to the F_0_, where recombination events never generate new ancestry breakpoints, we effectively compensate for this effect, at least as long as *E*[*x*(0)] ≈ *E*[*x*(1)]. This yields an expected mapping resolution of:

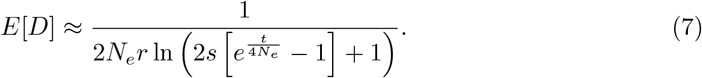

In the following, we will refer to Eq. (4) as the “recursion” solution, while the approximation presented in Eq. (7) will be referred to as the “integration” solution.

### Limiting cases

We now want to take a closer look at the expected mapping resolution derived in Eq. (7) and discuss how it relates to the result from the infinite population model. First, as we already mentioned above, our approach relies on the assumption that *t* ≪ 4*N*_*e*_, as drift would otherwise be strong and heterozygosity would be expected to decay noticeably over the course of the experiment. This assumption specifies a regime where the probability that a given pair of lineages coalesce over the course of the experiment is still small (since the expected time to pairwise coalescence should be 2*N*_*e*_ generations). Given *t* ≪ 4*N*_*e*_, we can perform a Taylor series approximation to the exponential in Eq. (7):

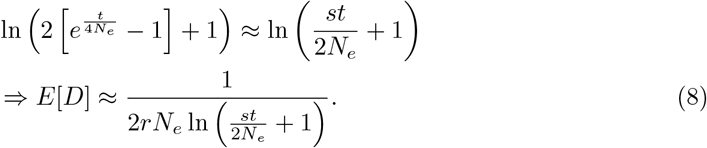

This approximation allows us to better understand how the infinite and finite population models differ from each other. In the infinite population model, mapping resolution was simply inversely proportional to the product of recombination rate (*r*), sample size (*s*), and number of generations (*t*) of the experiment. In the finite population model, mapping resolution is still inversely proportional to the recombination rate, but the effects of sample size and experiment length are now attenuated by a logarithm. Consequently, increasing those parameters is no longer expected to improve mapping resolution as effectively as in the infinite population model. We further note that sample size and generations enter Eq. (8) only in terms of the product *s* × *t*. Varying each of these two parameters by the same factor is therefore expected to produce a similar impact on the expected mapping resolution (as long as *t* ≪ 4*N*_*e*_ still holds). In practice, this means that running an experiment twice as long, for instance, should yield the same benefit as doubling the sample size.

Eq. (8) also reveals where the effects of a finite population start to become substantial. When *st* ≪ 2*N*_*e*_, we can further approximate

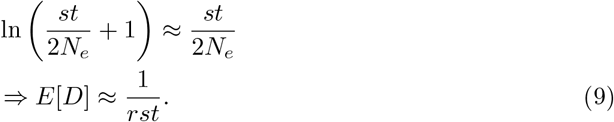

Thus, the infinite and finite population models converge in this regime. The two models will increasingly diverge from each other as *st* becomes of the same magnitude as 2*N*_*e*_. The condition *st* ≪ 2*N*_*e*_ should typically be much stricter than *t* ≪ 4*N*_*e*_, our essential assumption for the finite population model, unless sample size is very small. The former effectively assumes that there are only very few coalescence events among the genealogy of all sampled chromosomes, whereas the latter only assumed that coalescence was unlikely between any two sampled chromosomes.

Figure 4 illustrates the behavior of our analytical solutions for the finite and infinite population models as a function of the product *st*, and, in the finite population model, for different values of *N*_*e*_. As predicted, both models converge when *st* ≪ 4*N*_*e*_. Compared to the infinite population model, increasing *st* provides only diminishing returns for improving mapping resolution in the finite population model. Lower *N*_*e*_ values generally decrease mapping resolution.

**Figure 4:**
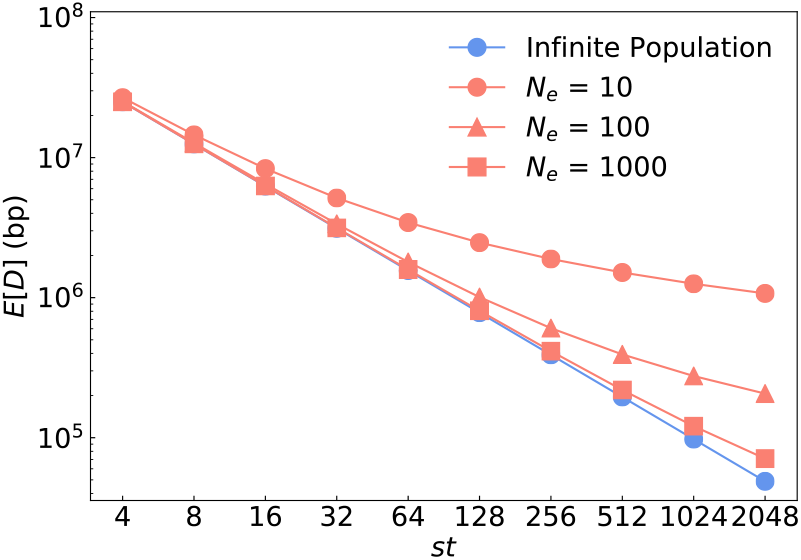
Analytical solutions for the infinite and final population models. Blue dots show the prediction by the infinite population model according to Eq. 1; red dots show the prediction by the finite population model according to Eq. 6 for 3 different values of *Ne*. Recombination rate was set to *r* = 10^−8^ per bp. To vary *st* in these equations, we fixed *t* = 2 and then varied *s* from 2 to 1024. The infinite and finite population models converge as *st* becomes much larger than 2*Ne*, as predicted by our theory.

### Numerical Validation

To evaluate the accuracy of our analytical results, we conducted individual-based simulations of a BSA experiment (see Methods). Specifically, we modeled an experimental setup as described in Figure 1A, assuming a trait that is determined by a single QTL located on a 100 Mbp-long chromosome. We assumed a uniform recombination rate of *r* = 10^−8^ per bp and generation (i.e., 1 cM/Mbp), which we did not vary in our simulations because mapping resolution should always be inversely proportional to *r*. The parameters we did vary were the sample size (*s*), the effective population size (*N*_*e*_), and the number of generations of interbreeding (*t*).

Figure 5 shows the comparisons between these simulations and our analytical results given by Equations (1), (4), and (7) over a broad range of parameter values (*N*_*e*_ varying between 10 - 1000, *s* varying between 2 - 1024, and *t* varying between 2 - 20). For each parameter setting, we estimated *D* over 5000 simulations. The resulting distributions are shown by box-and-whisker plots. Note that these distributions tend to have rather pronounced positive skews, such that their means tend to be much larger than their medians. Our analytical results are given in the form of expectation values for *D*, and thus need to be compared to the mean values of the simulation data, not the medians.

**Figure 5:**
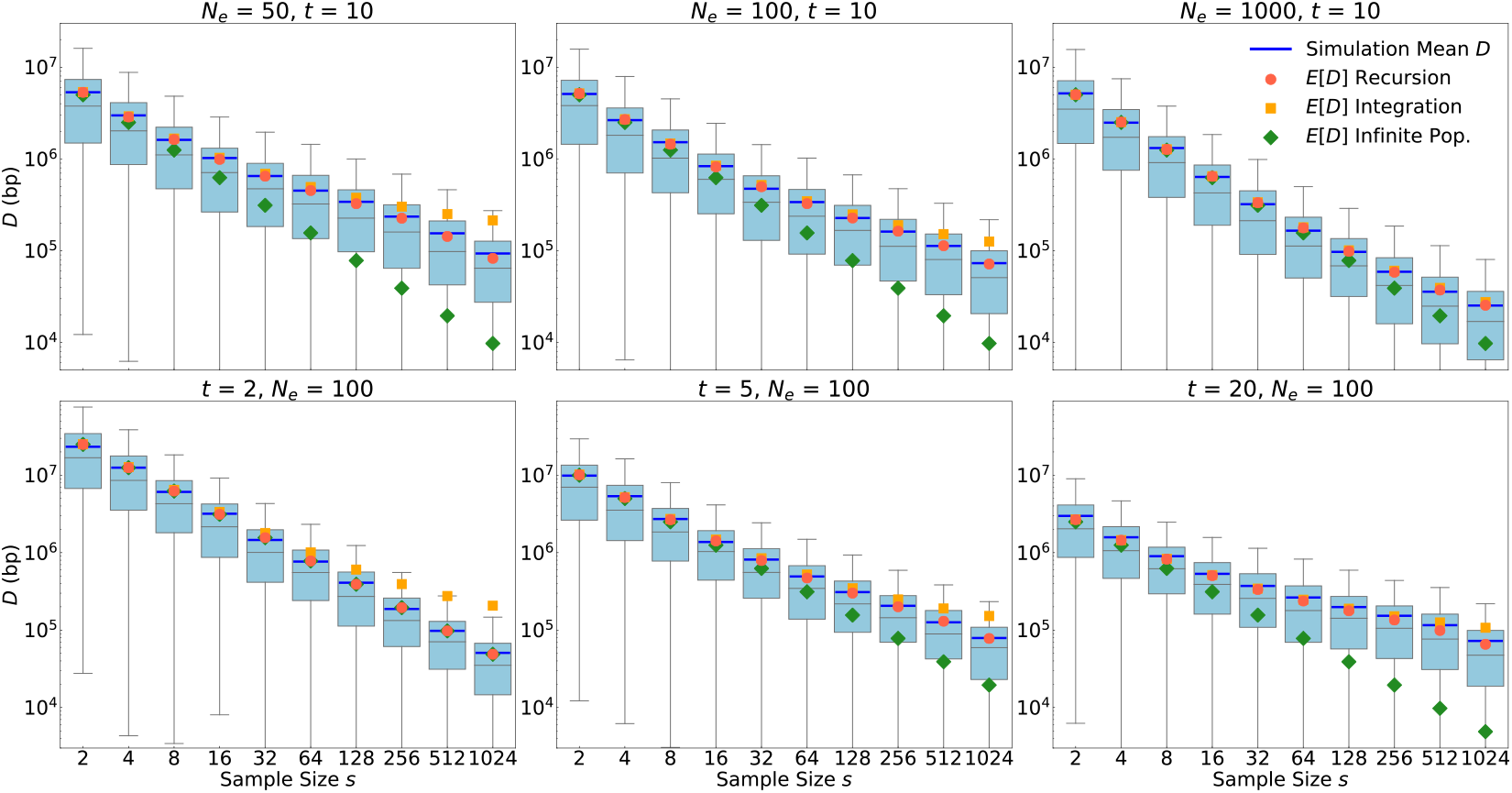
Comparison between analytical results and simulations. A single QTL located on a 100 Mb chromosome with a uniform recombination rate of 1 cM/Mb was modeled. Box plots show the distribution (quartiles) of *D* estimated over 5000 simulation runs for each given parameter setting. The top/bottom whiskers represent the highest/lowest datum within the 1.5 interquartile range of the upper/lower quartile. Blue lines show the means of the data, which tend to be much larger than the medians. Symbols show the expected resolutions for the infinite (green) and finite (red/orange) population models according to Equations (1) and (4/7), respectively. Green dots are difficult to see in the lower left panel because they are almost completely overlaid by the red dots. Note that these analytical predictions should be compared to the means of the simulations (blue lines), not the medians. In the top row, we varied *Ne* from 50 to 1000 while keeping *t* = 10 constant. In the bottom row, we varied *t* from 2 to 20 while keeping *Ne* = 100 constant.

The simulation results show excellent agreement with our recursion solution for the finite population model provided in Eq. (4). As already discussed above, the solution from the infinite population model provided in Eq. (1) constitutes an upper bound for the maximally achievable mapping resolution. Consistent with analytical predictions, the finite and infinite models converge when *st* ≪ 4*N*_*e*_, and the infinite population model increasingly overestimates mapping resolution as the condition *st* ≪ 4*N*_*e*_ is increasingly violated.

The integration approximation of the finite population model we derived in Eq. (7) generally works well for small and moderate sample sizes, but tends to underestimate mapping resolution when *s* approaches *N*_*e*_ in magnitude. It also breaks down when *t* is very small (as can be seen in the lower left panel for *t* = 2, where the integration approximation actually becomes less accurate than the infinite population model). This is a consequence of the use of a continuous integration in Eq. (6), which underestimates the total tree length when the number of discrete generations is small. However, in this small *t* regime, the recursion solution, which is most accurate, can also be easily evaluated due to the need for only few recursion steps.

### Extension to alternative crossing schemes

Our analytical approach for calculating *E*[*D*] is straightforward to extend to variations of the experimental design, such as alternative crossing schemes. The key parameters that need to be ascertained for a given design are the rate *ρ*(*i*) at which new ancestry breakpoints are generated per bp in the gametes that will make up the individuals in generation *i*, together with *E*[*x*(*i*)], the expected number of ancestral lineages present in the sample’s genealogy in that generation. The expected mapping resolution is then given by a direct generalization of Eq. (3):

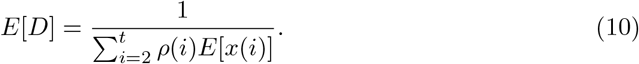

In the standard BSA design, we had *ρ*(2) = *r* and *ρ*(*i >* 2) = *r/*2, with *E*[*x*(*i*)] calculated recursively by Eq. (2) using the coalescence effective size *N*_*e*_ of the interbreeding population.

In the following, we will illustrate how this approach can be applied to two proposed modifications of the standard BSA design. The first approach is introgression mapping (IM), illustrated in Figure 6 (left). Here, *AA* homozygotes are selected in every even generation of the experiment (the approach thus relies on our ability to do so effectively). These individuals are then backcrossed to the *aa* parental strain. The resulting offspring are interbred without selection in every odd generation, after which the cycle starts anew. At the end of the experiment, *s* individuals of genotype *AA* are selected and sequenced. Their genomes should then resemble the *AA* strain across a genomic region that surrounds the causal QTL, while resembling the *aa* strain throughout the rest of the genome.

**Figure 6:**
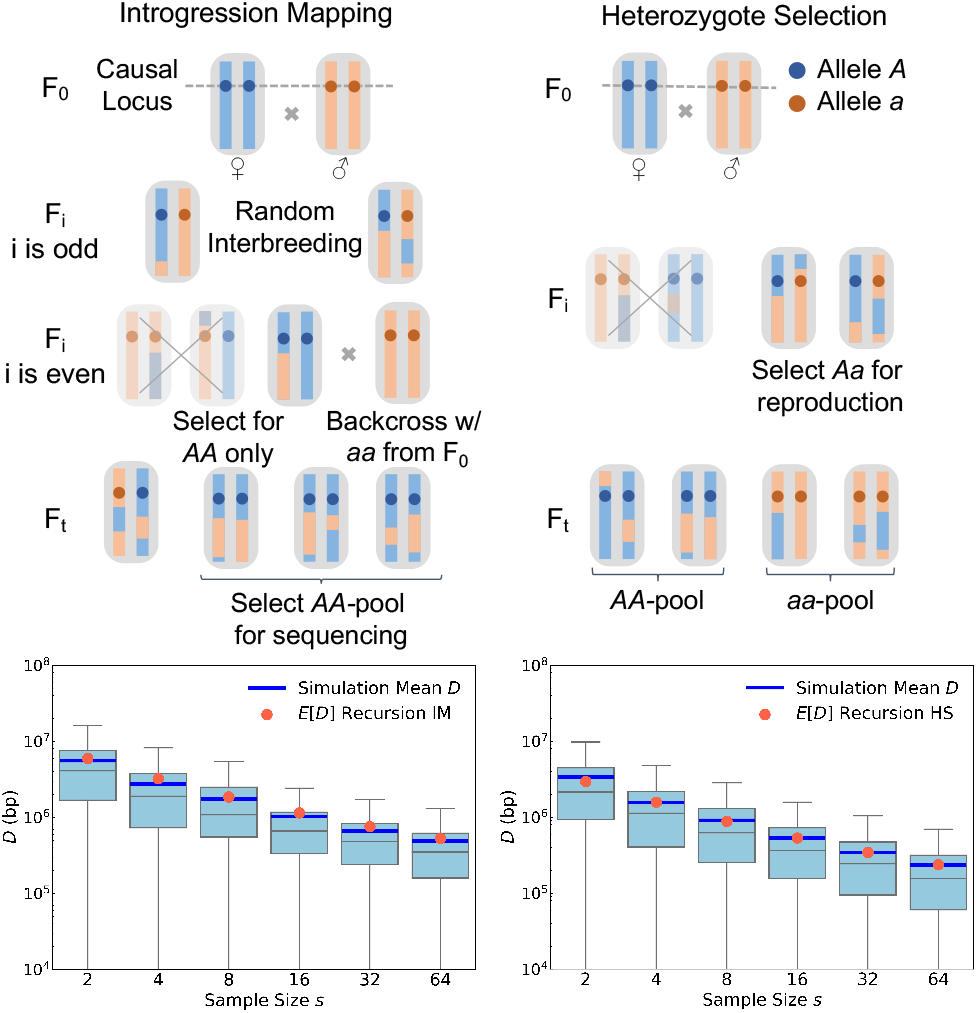
Extension of our theory to two different crossing schemes. In the introgression mapping (IM) scheme shown in the left panels, *AA* homozygotes are selected in even generations and then backcrossed to the *aa* founding strain. The resulting offspring are interbred without selection in odd generations. At the end of the experiment, *AA* homozygotes are sequenced. The bottom-left panel shows that the distributions of *D* values in simulated IM experiments conform well to our analytical predictions (see Methods). The heterozygote selection (HS) scheme shown in the right panels is similar to the standard BSA design, except that only *Aa* heterozygotes are allowed to reproduce in every generation. Our theory again accurately predicts the expected mapping resolution under this design (bottom-right panel).

During the odd generations of an IM experiment, all individuals should be *Aa* heterozygotes at the QTL, and thus carry one red and one blue ancestry segment across some region surrounding it. These segments will become shorter and shorter due to recombination events as the experiment progresses. The rate at which new ancestry breakpoints are created close to the QTL, during the odd generations, should therefore be twice that of a standard BSA experiment, while it will be zero in all even generations (when all surviving individuals will be *AA* homozygotes at the QTL). When averaged over the whole experiment, new ancestry breakpoints in the vicinity of the QTL should hence arise at a rate of *r/*2, similar to the BSA design.

However, due to the selection step for *AA* homozygotes in the even generations, the coalescence rate in the IM design should be higher as compared to a standard BSA design with an interbreeding population of comparable size, given that only 1*/*4 of the population should be *AA* homozygotes. Thus, the value of *N*_*e*_ will need to be adjusted in Eq. (2). A reasonable approximation would be to use the harmonic mean between odd and even generations, yielding *N*_*e*_ = 0.4*N*′, where *N*′ is the coalescence effective population size of the interbreeding population in a scenario where no selection for homozygotes would be performed. Figure 6 confirms that this approach produces an accurate analytical prediction for the expected mapping resolution in an IM experiment.

This example illustrates how our theory can help evaluate the expected performance of alterations to an experimental design. For the IM design in particular, the fact that *ρ*(*i*) should be comparable to a standard BSA design when averaged over the entire experiment, while *N*_*e*_ should be smaller, suggests that an IM design should generally have lower resolution than BSA, confirming previous simulation results (Pool 2016). Yet, there may be other advantages of IM. For example, this design ensures that *A* and *a* alleles are kept at 50% frequency throughout the experiment, thereby eliminating any potential effects of drift or selection at the QTL that could exist in a standard BSA design.

The second alternative design we want to discuss is heterozygote selection (HS), illustrated in Figure 6 (right). In this approach, only *Aa* heterozygotes are selected for reproduction in every generation (again assuming that we can do so effectively). This should double the rate of ancestry breakpoint generation in the vicinity of the QTL as compared to a standard BSA design, so that *ρ*(*i*) = *r* for all generations *i* ≥ 2. However, the effective population size will again be reduced due to the selection step. Here, a reasonable approximation should be that *N*_*e*_ is about 1/2 of that in a standard BSA design with an interbreeding population of comparable size, given that about half of the population are expected to be *Aa* heterozygotes at any point. Our simulations confirm that this approach again produces an accurate analytical prediction for the expected mapping resolution in an HS experiment (Figure 6).

In principle, due to the higher rate of ancestry breakpoint generation, the HS design could therefore yield a mapping resolution up to two times better than a standard BSA design, as long as this is not outweighed by the concomitant reduction in *N*_*e*_. Note that, as with IM, the HS design maintains the frequency of *A* and *a* alleles at 50%.

## Methods

Simulations of BSA experiments were implemented in the SLiM (version 3.5) evolutionary simulation framework [13]. We modeled a single QTL located on a 100Mb chromosome. Each experiment was initialized with two homozygous parental strains (denoted as *AA* and *aa* strains). The F_1_ was always seeded with 1000 males from the *AA* strain and 1000 females from the *aa* strain. The population was then interbred over *t* discrete, non-overlapping generations, using SLiM’s default Wright-Fisher model without selection. While the total number of individuals was kept constant at 2000 in each generation, only *N*_*e*_ randomly chosen individuals were actually allowed to mate and reproduce in each generation. Recombination occurred at a uniform rate of *r* = 10^−8^ per bp (i.e., 1 cM/Mb) along the chromosome in all simulations.

For the comparisons of analytical vs. simulation results in the standard BSA design (Figure 5), as well as the HS scheme (Figure 6), we used SLiM’s tree-sequence recording feature [12] to track the ancestry at each position in each genome. This allowed us to directly identify ancestry breakpoints in the sampled chromosomes without having to model any marker SNPs for such inference.

For the simulations of the IM scheme (Figure 6), we modeled SNPs placed at equidistant intervals of 10 kb along the chromosome to differentiate ancestry from the two parental strains. While this approach only allows for indirect and approximate inference of ancestral breakpoint locations, it should not pose a limiting factor given that the mapping resolution was several orders of magnitude larger than the distance between marker SNPs.

For the simulation of a short-read PoolSeq experiment (Figure 7), we used marker SNPs placed at equidistant intervals of 1 kb along the chromosome. Here, we assumed that every SNP provided an independent locus where a new set of *C* chromosomes was genotyped from the *s* chromosomes present in each sample. These chromosomes were chosen randomly with replacement. Thus, we effectively assumed a read length of less than 1 kb.

**Figure 7:**
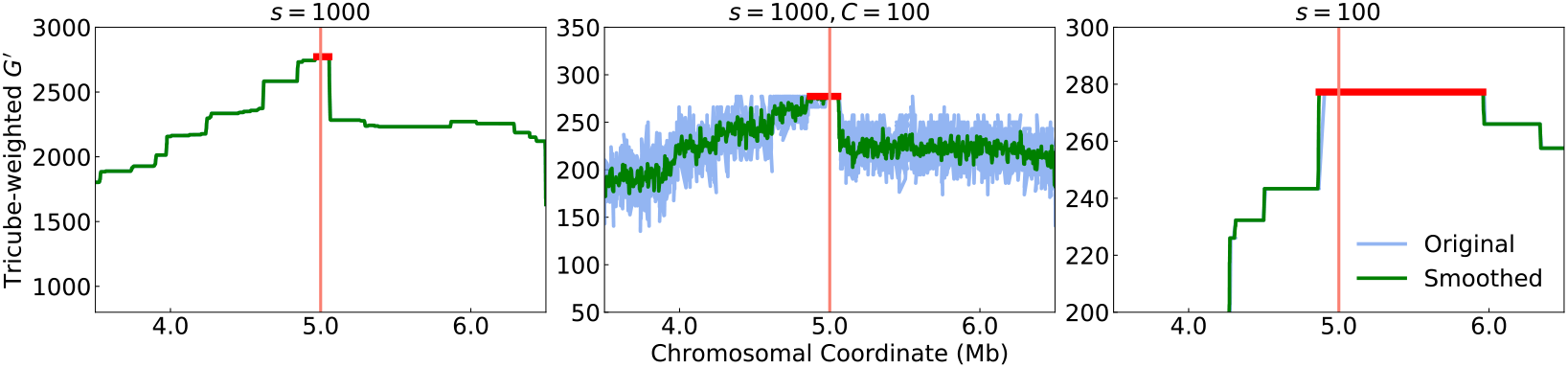
The impact of sample size versus coverage in a BSA experiment with short-read sequencing. All simulations modeled a single QTL (vertical red line) at the center of a 10 Mb-long chromosome in a BSA experiment with *Ne* = 100 and *t* = 10. The outside plots show two example simulation runs of experiments with sample sizes *s* = 1000 (left) and *s* = 100 (right), where we assumed perfect sequencing data for every genome in the two pools. The middle plot shows a simulation with sample size *s* = 1000 as before, but here with the addition of simulated short-read sequencing at only *C* = 100× coverage in each pool. Blue lines show tricube-weighted *G*′ statistics, while green lines show a smoothed version of that statistic. The red horizontal bars indicate the size of the peak around the causal locus in which *G*′ is maximal, thereby determining mapping resolution. As expected, larger sample size yields better mapping resolution. Remarkably, the low-coverage experiment yields a similar resolution to the full-coverage experiment, despite having less than 10% of genomes, on average, actually genotyped at each locus.

The calculation of *G* in Figures 1B and 7 followed the procedure illustrated in [21]. Smoothed curves were obtained by a weighted sum of all SNPs within the window bracketing the focal SNP, where the weight of each SNP was obtained by a Nadaraya-Watson kernel regression [35].

The SLiM model for simulations, Python scripts for data analysis, and other related files are available at: https://github.com/runxi-shen/Predict-Genomic-Resolution-of-BSA.

## Discussion

BSA has become an increasingly popular technique for mapping the genetic basis of phenotypic traits. Previous studies have used simulations to study how the genomic resolution of BSA is affected by key experimental parameters such as sample size and number of generations of interbreeding [26]. However, a truly quantitative understanding has so far remained elusive. In this study, we were able to derive an analytic solution for the expected mapping resolution of a BSA experiment. We have further demonstrated how our framework can be easily extended to modifications of the experimental design, such as introgression mapping or selection for heterozygotes.

Our approach is based on the insight that the mapping resolution of a BSA experiment is ultimately limited by the length of the genomic region surrounding the QTL in which all sequences in each sampled pool still share the ancestry of the respective parental strain. This region is delimited by the two closest ancestry breakpoints observed upstream and downstream of the QTL. We model the occurrence of such breakpoints by a Poisson process along the chromosome, with its rate determined by two factors: the expected length of the sample’s genealogy at any given genomic position, and the expected rate at which new ancestry breakpoints were generated along this genealogy in the ancestors to the sample. Both factors combine to determine the expected mapping resolution according to Equation (10). Our solution sheds light on the possible avenues for improving mapping resolution. First, the rate of ancestry breakpoint generation could be increased. While this rate is obviously bounded by the recombination rate of the organism, only recombination events in individuals that carry ancestry segments from both parental strains at the crossover location actually generate new ancestry breakpoints. Thus, one could seek to increase the frequency of such individuals; this is the rationale behind the “heterozyogte selection” strategy we discussed above. Second, the length of the sample’s genealogy could be increased. In principle, this could be achieved by using a larger sample size, including more generations of inbreeding, or achieving a lower coalescence rate during the experiment. Exactly how these parameters play out will depend on the specific experimental setup.

In Equation (7), we provided an approximate solution for the maximal mapping resolution of a standard BSA experiment. This solution requires specification of the coalescence effective population size (*N*_*e*_) of the interbreeding population that determines the pairwise coalescence rate in the genealogy of the sample. In practice, the value of *N*_*e*_ will typically be smaller than the actual number of individuals present in the interbreeding population, especially when there is high variance in offspring number among individuals [5]. While various methods have been developed for inferring *N*_*e*_ of experimental populations [2,16,19,36], such inference may be non-trivial, and it may thus be unclear how to choose the appropriate value for this parameter; at a minimum, however, the population size of the interbreeding population constitutes an upper bound for *N*_*e*_. Our analysis suggests that *N*_*e*_ should generally be kept as large as possible throughout the experiment to optimize mapping resolution.

The way in which we assess mapping resolution in our theory may not always correspond exactly to what determines the resolution in a real-world experiment. For example, our mathematical approach is based on the location of ancestry breakpoints in the sampled chromosomes, but such breakpoints are typically not directly observable. Instead, their location can only be inferred approximately through marker SNPs that allow one to distinguish ancestry from the two founding strains. The genomic density of such marker SNPs thus places a practical limit on the achievable mapping resolution; however, this should not be problematic as long as the average distance between differentiating sites remains short compared to the mapping resolution predicted by our theory. Another simplification is that we have assumed perfect sequencing, while any experiment will suffer from some level of sequencing errors. This will create “noise” in the summary statistics used for mapping such as *G*′.

Perhaps most importantly, the sequencing data in a BSA experiment will typically be comprised of short-read sequences from pooled samples [30], rather than individual genome sequences. Unless the sequencing coverage level substantially exceeds the sample size, the number and specific set of genomes that are actually sequenced at any given genomic position will thus vary along the genome. This raises an important practical question: is sample size or coverage (*C*) the more critical factor in limiting the resolution of a BSA experiment? Our theory makes a clear prediction. Since mapping resolution is ultimately limited by the ancestry breakpoints present in the sampled genomes, the size of the samples will be the primary factor. In larger samples, there is simply a better chance to capture more breakpoints that are closer to the QTL. By contrast, even for low coverage (*C* ≪ *s*), it is still possible to achieve a mapping resolution close to what would be predicted by our theory for the given sample size. This is due to the randomness in the pooled sequencing process. Consider, for example, two SNPs that are separated by a distance that is substantially larger than the typical read length. We will then likely find different sets of sampled chromosomes being sequenced at each. Thus, as we move away from the QTL, each such locus can be regarded as a new trial that provides another chance to observe a read with ancestry from the opposite strain, thereby allowing us to confine the location of the causal locus. As long as read length is much shorter than the expected mapping resolution, the large number of trials can thus make up for the limited number of chromosomes sequenced at each individual locus.

In Figure 7, we confirm this prediction for simulated BSA experiments, demonstrating that short-read sequencing at a coverage level ten times smaller than the sample size still achieves a mapping resolution similar to that achieved with full sequencing coverage of the entire sample. Interestingly, our theory suggests that substantially longer reads would actually perform worse in such situations, as this would increase the distance over which the set of sequenced genomes would remain correlated.

For the sake of mathematical tractability, we have focused on a rather simplistic model of a trait controlled by a single QTL. Our results should still hold for traits determined by multiple loci, as long as these loci are on separate chromosomes. However, when several QTL are located on the same chromosome, overlap between signals could become a problem [26]. Furthermore, it may no longer be possible to select for individuals that are homozygous for the genomic background of the respective parental strain at all QTL. In such cases, we expect that our theory can only provide a lower bound to the achievable mapping resolution. Our approach further assumed that recombination rate is uniform along the chromosome, but it would be rather straightforward to incorporate non-uniform recombination rates. In particular, recombination events would then need to be modeled by an inhomogeneous Poisson process in Eq. (3), so that *R*, and thus also *E*[*D*], would become a function of genomic position. Intuitively, it is clear that we expect higher mapping resolution in regions of higher recombination rate, and vice versa.

In the early days of molecular genetics, the precision one could hope to achieve in a mapping experiment was typically limited by the ability to genotype a sufficient number of individuals at a sufficiently dense set of marker loci. With the sequencing revolution, this constraint has fundamentally shifted. Today, it is often feasible to obtain whole-genome sequencing data for samples of several hundreds or even thousands of individuals. Consequently, it is becoming more relevant to understand which other factors fundamentally limit mapping resolution under a given experimental design. By providing an analytic solution to the expected mapping resolution of a BSA experiment, based on coalescence theory, we were able to shed light on how individual parameters combine, qualitatively and quantitatively, to place a fundamental limit on mapping resolution. We hope that these results can not only help scientists to set realistic expectations for the power of their planned experiments, but also to identify which strategies would allow them to optimize their study design most efficiently and economically. Finally, we hope that the conceptual approach that underlies our theory can be extended to other genetic mapping strategies.

